# Identification of circular RNAs in porcine sperm and their relation to sperm motility

**DOI:** 10.1101/608026

**Authors:** Marta Gòdia, Anna Castelló, Martina Rocco, Betlem Cabrera, Joan E. Rodríguez-Gil, Armand Sánchez, Alex Clop

**Affiliations:** Animal Genomics Group, Centre for Research in Agricultural Genomics (CRAG) CSIC-IRTA-UAB-UB, Campus UAB, Cerdanyola del Vallès (Barcelona), Catalonia, Spain; Unit of Animal Science, Department of Animal and Food Science, Autonomous University of Barcelona, Cerdanyola del Vallès (Barcelona), Catalonia, Spain; Unit of Animal Reproduction, Department of Animal Medicine and Surgery, Autonomous University of Barcelona, Cerdanyola del Vallès (Barcelona), Catalonia, Spain; Consejo Superior de Investigaciones Científicas (CSIC), Barcelona, Catalonia, Spain

**Keywords:** sperm, circular RNA (circRNA), RNA-seq, miRNA, sperm motility, sperm quality, swine, male infertility

## Abstract

Circular RNAs (circRNAs) are emerging as a novel class of noncoding RNAs which potential role as gene regulators is quickly gaining interest. Although circRNAs have been studied in several tissues and cell types across several animal species, the characterization of the circRNAome in ejaculated sperm remains unexplored. In this study, we profiled the sperm circRNA catalogue in 40 boar samples. A complex population of 1,598 circRNAs was shared in at least 30 samples. The predicted circRNAs presented in general low abundances and were highly tissue-specific. Circa 80% of the circRNAs identified in the boar sperm were reported as novel. We also constructed a circRNA-miRNA interaction network based on experimental and predictive microRNA (miRNA) binding sites. Moreover, we found significant correlation between the abundance of some circRNAs and sperm motility parameters and confirmed two of these correlations by RT-qPCR in 36 samples with extreme sperm quality. Our study provides a thorough characterization of a novel cell type for the circRNA encyclopedia collection and suggests that circRNAs are potential noninvasive biomarkers for male sperm quality and thus, might also hold potential to predict male fertility.

## Introduction

Food scarcity is a growing concern for our society due to human population growth and climate change. Swine and poultry are the main sources of meat worldwide and large efforts are being taken to increase their production in a sustainable manner. In this sense, reproductive efficiency is one of the key aspects for the sustainability of animal breeding [1]. In addition, due to their similarity in genome sequence, anatomy and physiology, swine is quickly becoming an important model for human bio-medical research [2]. In humans, infertility is an increasing problem in contemporary society, affecting one in twenty males [3], and has become a subject of bio-medical research in swine [4–6]. Unlike in humans, where fertility data is based on few records per person, the swine industry has begun to record fertility traits each time a male is used for artificial insemination (AI). In pig intensive production systems, during its productive life, an AI boar typically inseminates around 1,500 sows (Balasch, personal communication). Moreover, the porcine ejaculates are routinely evaluated in the AI centers after every extraction. Thus, large datasets of the reproductive ability of these boars is becoming available. Taken all together, swine becomes a good animal model to study semen quality and male fertility.

Sperm motility and kinetic parameters provide an objective and reproducible measurement of semen quality that is automatically assessed by the computer-assisted semen analysis (CASA) system. CASA records the percentage of total motile sperm, curvilinear velocity (VCL), straight line velocity (VSL) and velocity of the sperm cells (VAP) and head frequency based on trajectories of motile sperm. This approach has been commonly used to assess semen quality in animal breeding strategies prior to AI in cattle [7], horse [8] and swine [9–11] where significant correlations between sperm motility and fertility have been found. In humans, this technique is also applied to estimate the *in vitro* fertilizing potential of the ejaculates used in assisted reproductive treatments [12–14].

Multiple research efforts have demonstrated that sperm quality parameters and fertility outcomes are related to the presence or absence of sperm RNAs. Different studies have provided evidence that the absence or deregulation of certain RNAs is associated to infertility and/or sperm motility as in human [15], mice [16] and cattle [17]. Studies have focused their research on messenger RNAs (mRNAs) and on different classes of non-coding RNAs. In humans, several microRNAs (miRNAs) have been found to be dysregulated in infertile patients [18]. Similarly in bulls, Capra *et al*. found differential abundances of some miRNAs and transference RNAs (tRNAs) between high and low motility sperm populations [19].

Circular RNAs (circRNAs) are a novel class of non-coding RNAs with a closed loop structure, mainly formed through pre-mRNA back splicing event [20]. CircRNAs are highly stable *in vivo* in comparison to their mRNA linear counterparts, because of their circular structure, which confers protection against exonucleases, ribozymes, antisense RNAs and small-interfering RNAs [21, 22]. Expanding views on their biogenesis and function suggests that circRNAs can directly regulate the abundance of their cognate mRNA and can also act as miRNA sponges [23], sequestering these miRNAs and thus impeding their post-transcriptional inhibitory roles on target mRNAs [23]. circRNAs have been identified across several species and tissues including the fruitfly [24], human [25, 26], mice [26, 27] and swine [28, 29]. These studies revealed species-, developmental- and tissue-specific expression patterns.

During the last few years, there has been a considerable interest in the potential use of circRNAs as biomarkers for health. Several studies have identified circRNAs which abundances were associated to cancer, aneurysms, hypertension, heart failure, diabetes and arthritis, among others [reviewed in 30]. In addition, there are few reports characterizing circRNAs in reproductive organs and cell types including oocytes, embryo, placental tissue, granulosa cells, immature spermatogenic cells and testis [reviewed in 31]. Up to date, a small number of studies have assessed the circRNA predictive potential for reproduction outcomes. For example, Chang and colleagues identified circRNAs associated to embryo quality and suggested their role as potential predictors of live birth [32] and Qian *et al*. identified differentially expressed circRNAs in the placenta of pregnant woman affected with preeclampsia [33].

Our group has recently carried a thorough porcine sperm transcriptome analysis [34] and the data, in line with previous studies [reviewed in 35], showed that the majority of the transcripts are highly fragmented and with low abundances. Considering their stability and abundance, circRNAs could hold an important potential as reliable biomarkers for sperm quality and fertility traits. Here, we have characterized the circRNA repertoire of the porcine spermatozoa, assessed their potential role as miRNA sponges and investigated the relationship between their abundance levels and sperm motility.

## Results

### Characterization of the sperm circular RNA repertoire

1,598 potential circRNAs were present and shared in at least 30 of the 40 ejaculates (Supplementary Table 1). The majority of the circRNA species were derived from exonic regions (CDS, 3’ and 5’ UTR) (82.1%), while only 13.5% and 4.4% circRNAs originated from intergenic and intronic segments, respectively (Figure 1A). Among the exonic circRNAs, most were conformed by less than 4 exons (81.0%), and just a few (14 circRNAs) contained 10 or more exons (Figure 1B). In addition, most exonic circRNAs (76.9%) were less than 400 bp long (Figure 1C). RNA abundance across the different circRNAs types was low, with a range between 0.19 and 136.4 CPMs and mean and median values of 2.42 and 0.89 CPMs, respectively (Supplementary Table 1). The top 20 most abundant circRNAs (Table 1) involved 15 coding genes and one annotated long non-coding RNA. These genes included *CEP63, ATP6V0A2, PPA2, PAIP2* and *PAXIP1*, which have been linked to sperm related traits and male fertility.

**Figure 1.**
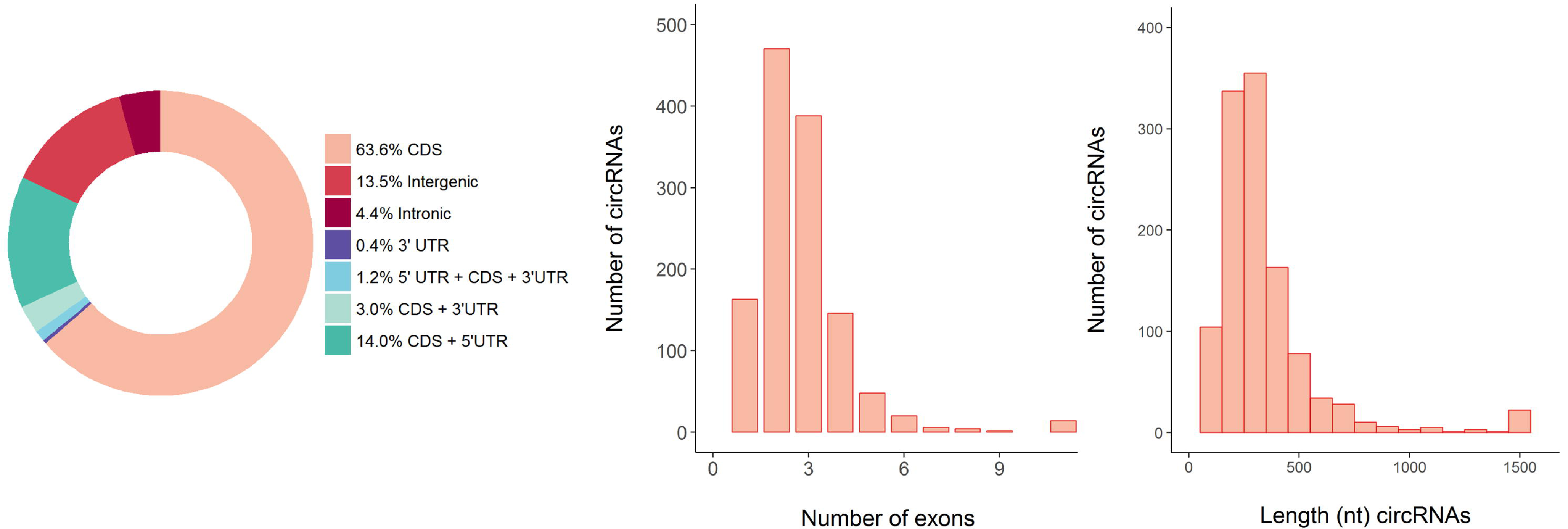
Characteristics of the genomic features of the sperm circRNAs identified in the boar sperm. **A.** Distribution of genomic location (CDS, intergenic, intronic, 3’ UTR or 5’UTR) of the 1,598 circRNAs identified in boar sperm. **B.** Distribution of the number of exons forming the exonic circRNAs. **C.** Distribution of the nucleotide length of the exonic circRNAs

**Table 1.** Top 20 most abundant circRNAs in swine sperm. circRNA genomic coordinates are indicated in the format chromosome:start_position..end_position. Mean and SD: standard deviation, are in CPM (Counts Per Million).

| circRNA name | circRNA genomic coordinates | Mean | SD | circRNA type | Ensembl ID | Host gene symbol |
| --- | --- | --- | --- | --- | --- | --- |
| ssc_circ_0493 | 13:75694364..75694548 | 136.4 | 80.7 | intronic | ENSSSCG00000011645 | *CEP63* |
| ssc_circ_0220 | 11:17636879..17653264 | 128.2 | 52.5 | intergenic |  |  |
| ssc_circ_0475 | 13:61967873..61969753 | 100.2 | 56.8 | lincRNA | ENSSSCG00000035295 |  |
| ssc_circ_1097 | 4:590305..590427 | 75.8 | 51.9 | CDS | ENSSSCG00000005916 | *WDR97* |
| ssc_circ_1141 | 5:22008783..22009750 | 75.0 | 35.4 | CDS | ENSSSCG00000000406 | *PTGES3* |
| ssc_circ_1062 | 4:130322062..130339222 | 62.2 | 33.1 | CDS | ENSSSCG00000006939 | *ZNHIT6* |
| ssc_circ_0537 | 14:29281863..29290551 | 59.9 | 34.8 | CDS | ENSSSCG00000009766 | *ATP6V0A2* |
| ssc_circ_0777 | 18:14243637..14259662 | 47.3 | 23.9 | CDS | ENSSSCG00000016529 | *AGBL3* |
| ssc_circ_0860 | 2:3096936..3098647 | 45.1 | 35.6 | CDS | ENSSSCG00000037451 | *PPFIA1* |
| ssc_circ_0805 | 18:54260018..54270584 | 40.0 | 52.3 | CDS | ENSSSCG00000035581 | *SUGCT* |
| ssc_circ_0132 | 1:641933..653683 | 33.2 | 19.1 | CDS | ENSSSCG00000004008 | *WDR27* |
| ssc_circ_1452 | 8:56584279..56590933 | 28.7 | 21.9 | intergenic |  |  |
| ssc_circ_1575 | AEMK02000682.1:1719140..1720285 | 28.4 | 24.6 | CDS | ENSSSCG00000005753 | *CAMSAP1* |
| ssc_circ_1413 | 8:116275909..116294261 | 26.9 | 15.5 | CDS | ENSSSCG00000022788 | *PPA2* |
| ssc_circ_0895 | 2:64840342..64840811 | 25.8 | 17.5 | 5' UTR | ENSSSCG00000013776 | *DDX39A* |
| ssc_circ_0102 | 1:264102211..264103554 | 24.3 | 22.1 | intronic | ENSSSCG00000005582 | *STRBP* |
| ssc_circ_0363 | 13:119893907..119895097 | 24.2 | 14.9 | inetergenic |  |  |
| ssc_circ_1101 | 4:6367950..6392035 | 22.6 | 28.8 | CDS | ENSSSCG00000005941 | *KHDRBS3* |
| ssc_circ_0839 | 2:141254413..141254577 | 21.8 | 20.9 | CDS + 5' UTR | ENSSSCG00000026606 | *PAIP2* |
| ssc_circ_0795 | 18:3101570..3102902 | 21.7 | 9.0 | CDS | ENSSSCG00000025221 | *PAXIP1* |

Only a small fraction of genes produce more than one circRNA. These are considered hotspot circRNAs genes. The circRNAs from hotspot genes are often produced from different back-splicing events of one exon with several others [29]. We detected 12 genes with 5 or more circRNA isoforms (Table 2). Some of these genes, namely *TESK2, SPATA19, PTK2* and *SLC5A10* are related to sperm function and fertility.

**Table 2.** circRNA hotspot genes in swine sperm. 12 genes presented 5 or more different exonic circRNAs.

| Gene symbol | Ensembl ID | Number of circRNAs |
| --- | --- | --- |
| *DENND1B* | ENSSSCG00000010900 | 8 |
| *DENND1A* | ENSSSCG00000005585 | 7 |
| *TESK2* | ENSSSCG00000003917 | 6 |
| *UIMC1* | ENSSSCG00000022508 | 6 |
| *ARMC9* | ENSSSCG00000023994 | 5 |
| *CAMSAP1* | ENSSSCG00000005753 | 5 |
| *KDM5B* | ENSSSCG00000010928 | 5 |
| *PTK2* | ENSSSCG00000038397 | 5 |
| *RPS6KC1* | ENSSSCG00000015586 | 5 |
| *SPATA19* | ENSSSCG00000025612 | 5 |
| *SLC5A10* | ENSSSCG00000018049 | 5 |
| *WNK1* | ENSSSCG00000000753 | 5 |

### Sperm circRNAs might be involved in epigenetic regulation and spermatogenesis

We analyzed the potential roles of the boar sperm circRNAs under the assumption that their function is related to their known mRNA counterpart. Gene Ontology (GO) analysis of the circRNAs host genes revealed an enrichment for epigenetic functions including histone modification (q-val: 5.52 × 10^−6^), histone H3-K36 methylation (q-val: 8.65 × 10^−3^) and chromatin organization (q-val: 2.16 × 10^−8^) (Supplementary Table 2). We also identified significant ontologies in spermatogenesis (q-val: 5.81 × 10^−4^), cilium assembly (q-val: 4.15 × 10^−3^) and developmental process (q-val: 1.33 × 10^−2^), among others (Supplementary Table 2).

### The boar sperm have a highly specific circRNAome

We compared our circRNA catalogue with equivalent available datasets [36] from other studies in human (including different brain sections tissues and several cell lines) [26, 37–40], mice (several brain segments, cell types and embryonic stem cells) [26, 37] and swine (lung, skeletal muscle, fat, heart, liver, spleen, kidney, ovarium, testis and 5 brain sections) [28, 29]. Twenty-four % and 11.3% of the boar circRNAs had potential human and mouse orthologs, respectively (Table 3). On the other hand, 20.3% of the porcine sperm circRNAs were also present in other porcine tissues (Table 3). The tissues showing higher overlap with sperm were testes (11.6%) and cortex (11.3%). All the other tissues were clearly less concordant (Table 3). By comparing this in the opposite direction, the proportion of the circRNAs already annotated in any pig tissue that were also present in our pig sperm list was 4.9%. This value was 12 times lower (0.4%) when evaluating the human circRNA catalog (Table 3).

**Table 3.** Concordance between the boar sperm circRNAs and other tissues. Number of circRNAs according to Sscrofa11.1 liftover, in human, mice and swine (detailed by tissue). The column % swine sperm shows the proportion of swine sperm circRNAs that were identified in the given species and tissues. Column % of the total circRNAs lists the proportion of circRNAs from that tissue or species that found a homolog in the boar sperm.

| Species / Tissues | Number of circRNAs | % swine sperm | % of the total circRNAs |
| --- | --- | --- | --- |
| Human | 90,067 | 24.0% | 0.4% |
| Mice | 15,498 | 11.3% | 1.2% |
| Swine | 6,663 | 20.3% | 4.9% |
| Basal ganglia | 456 | 2.6% | 9.2% |
| Brain stem | 820 | 5.3% | 10.2% |
| Cerebellum | 1,061 | 5.6% | 8.4% |
| Cortex | 2,163 | 11.3% | 8.4% |
| Hippocampus | 549 | 3.2% | 9.3% |
| Fat | 494 | 2.4% | 7.7% |
| Heart | 539 | 2.2% | 6.5% |
| Kidney | 469 | 1.9% | 6.4% |
| Liver | 353 | 2.1% | 9.3% |
| Lung | 683 | 2.7% | 6.3% |
| Muscle | 532 | 2.6% | 7.9% |
| Ovarium | 652 | 3.8% | 9.4% |
| Spleen | 241 | 1.4% | 9.1% |
| Testes | 2,685 | 11.6% | 6.9% |

### Sperm circRNAs do not follow an age-dependent pattern

We assessed whether sperm circRNAs depicted an age-accumulating profile as has been previously observed in rat testes [41] and rat and mouse brain tissues [41, 42]. All the boars from our dataset were mature (≥ 8 months old) with ages ranging between 9 and 54 months of age. Sexual maturity in boars is a process that starts at the age of 8 months and finalizes at the age of 2 years [43]. Thus, we divided the samples in those coming from boars approaching sexual maturity with ages below 2 years old and those produced by mature pigs with ages above 2 years old. There was no significant difference in the number of circRNAs identified (P-value: 0.68, Wilcoxon rank sum test) nor in their RNA abundance (P-value: 0.948) between the two groups. We repeated the analysis considering only extreme ages: young (N=4; between 8.6 and 9.2 month old) and mature (N=4; 29.9 to 54.6 month old). Again, there was no difference in the number of circRNAs identified (P-value: 0.89) or in their abundance (P-value: 0.2).

### circRNA-miRNA interaction network

To characterize the potential role of the circRNAs as miRNA sponges, we built a co-expression network including the abundances of the 95 miRNAs and the 1,598 circRNAs identified in the short and total RNA-seq datasets. The association analysis resulted in 2,323 significant interactions between the 95 miRNAs and 564 circRNAs. On the other hand, the *in silico* prediction of miRNA targets in the circRNA sequences, based on sequence complementary, yielded 4,987 potential targets involving all the miRNAs and 1,103 circRNAs. To reduce the proportion of false-positives, only the 70 interactions (from 31 miRNAs and 56 circRNAs) that were found shared in both methods were used for network visualization (Figure 2). Most circRNAs (46) presented one miRNA target site with the exception of 10 circRNAs that harbored 2 or more potential targets (Figure 2). As the ssc_cic_08954 and ssc_circ_1454 which can potentially regulate, each, 4 distinct miRNAs (Figure 2). On the other hand, miR-26a and miR-28-5p were predicted to be regulated by 9 different circRNAs and 5 different circRNAs regulated miR140-3p and miR-423-5p (Figure 2).

**Figure 2.**
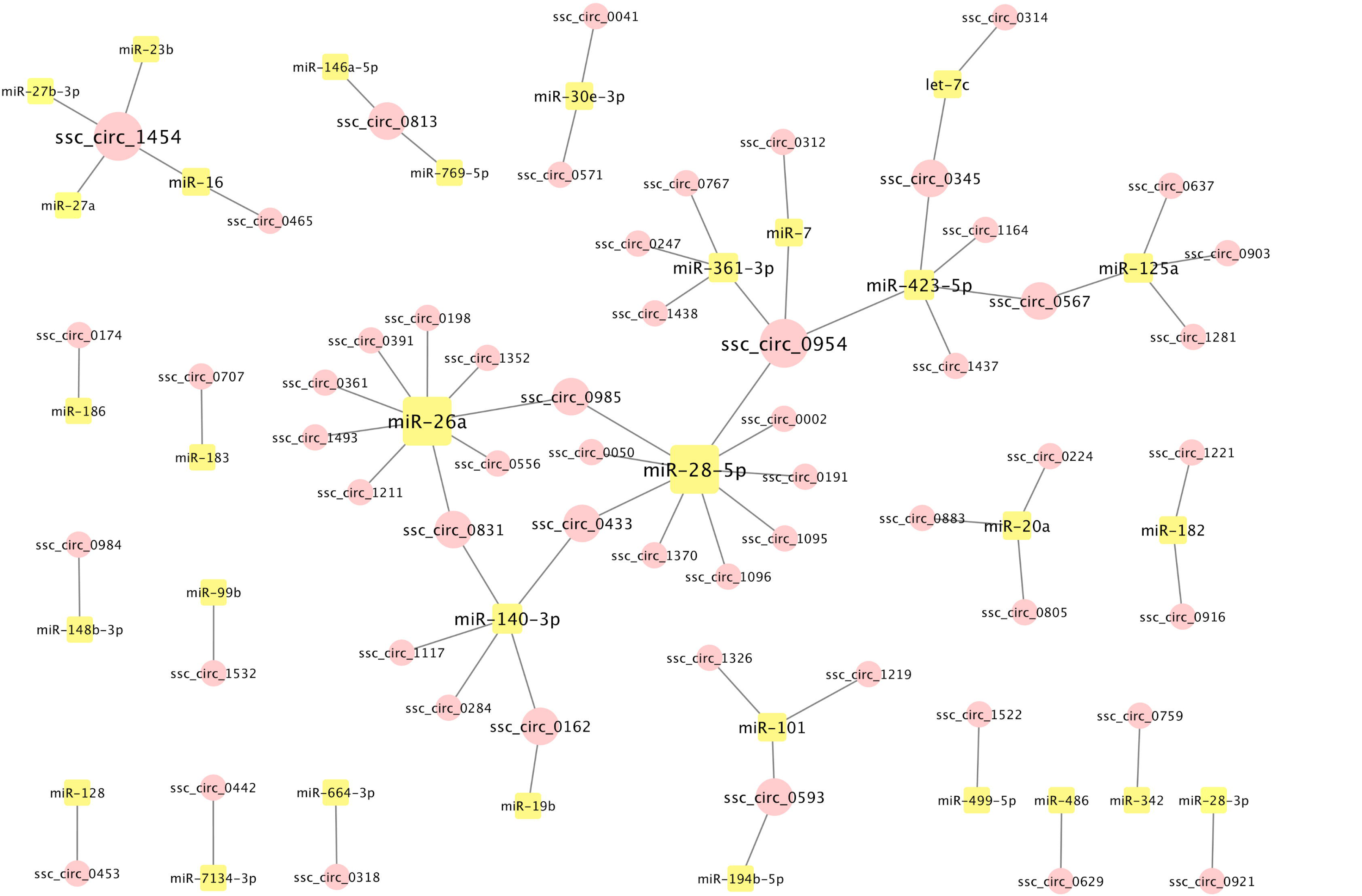
circRNA-miRNA interaction network. circRNA:miRNA relationships predicted by both experimental data and miRNA target prediction in circRNAs. Circular and square nodes represent circRNAs and miRNAs, respectively. The node and letter sizes indicate the number of significant correlations involving the node.

### Correlation of circRNAs with sperm motility and circRNA validation

179 circRNAs (from 26 intergenic regions, 10 intronic regions and 146 genes) showed significant associations between their abundance and different sperm motility parameters (Supplementary Table 3). More in detail, 28 (2 intergenic, 2 intronic and 24 protein coding genes), 94 (7 intergenic, 2 intronic and 81 genic), 35 (6 intergenic, 3 intronic and 26 genic) and 57 (11 intergenic, 3 intronic and 43 genic) circRNAs correlated with the percentage of motile cells, VCL, VAP and VSL, respectively (Supplementary Table 3).

To confirm the existence of these circRNAs and their phenotypic correlations we undertook a RT-qPCR and Sanger Sequencing approach. First, we randomly selected 2 circRNAs with different RNA abundance levels (75.0 and 10.6 CPM) to test whether we could confirm the bioinformatically predicted circRNAs. The chosen circRNAs were: ssc_circ_1141 (from *PTGES3*) with 75.0 CPM, and ssc_circ_0670 (from *BAZ2B*) with 10.6 CPM, both also detected in human [36] (hsa_circ_0008137 and hsa_circ_0002463, respectively) and in swine testes [28]. The PCR amplification of the 2 circRNAs resulted in a single electrophoretic band of the expected size (Supplementary Figure 1.A) and Sanger Sequencing confirmed the back-splice junction (Supplementary Figure 2.A-B). We, therefore, confirmed the validity of the RT-PCR and Sanger Sequencing approach to validate the presence of tested circRNAs and the existence of these two circRNAs in the boar sperm.

Then, we used the same approach to confirm the existence of 8 circRNAs selected for their significant correlation with at least one motility trait (Supplementary Table 3). These circRNAs were further selected based on the fact that (i) they were present at high abundances, or (ii) they showed significant correlation with at least 1 trait, or (iii) they had been previously identified in pig [28], human [36] or mice [36]. PCR amplification of 6 of the 8 circRNAs resulted in a band of the expected size (Supplementary Figure 1B) and the Sanger Sequencing validated the expected back-splice junction (Figure 3.A-C; Supplementary Figure 2.C-E). ssc_circ_0839 from *PAIP2* did not amplify (data not shown) and ssc_circ_0118 from *PDE10A* (also identified in pig testes and in human as hsa_circ_0078638) displayed 2 amplification bands (Supplementary Figure 1B). These two circRNAs were discarded for further analysis. Thus, we also confirmed the existence of these 6 circRNAs.

**Figure 3.**
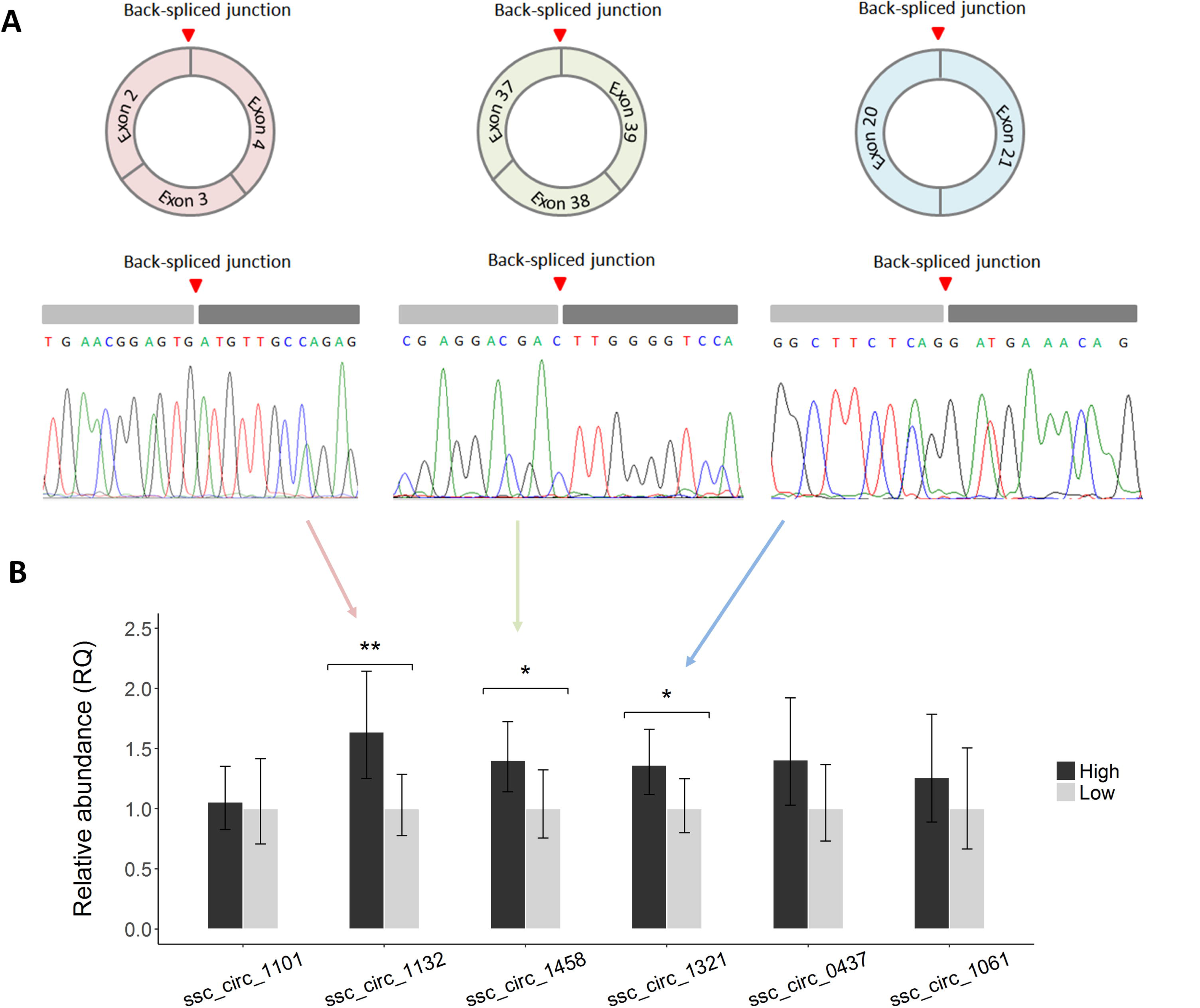
Validation of circRNAs that displayed RNA-seq abundance correlation with sperm motility. **A.** Sanger sequencing validation of the circRNAs black splice junction for ssc_circ_1132 from LIN7A, ssc_circ_1458 from LRBA and ssc_circ_1321 from PAPOLA. **B.** Relative abundance of six circRNAs in samples with extreme and divergent motility values (18 high and 18 low) obtained by RT-qPCR. The RNA-seq association between ssc_circ_1458 and ssc_circ_1321 and sperm motility was validated by RT-qPCR.

The RT-qPCR levels of these 6 circRNAs were measured in 36 animals presenting extreme and opposite values of sperm motility (N=18 for each phenotypic distribution tail) from a bank of 300 phenotyped boar ejaculates. None of the 36 samples was included in the RNA-seq study. The two sample groups displayed significant phenotypic differences for all the studied traits: percentage of motile cells (P-value: 2.65 × 10^−9^, Wilcoxon rank sum test), VCL (P-value: 2.20 × 10^−10^), VSL (P-value: 3.22 × 10^−7^) and VAP (P-value: 3.22 × 10^−7^). The RT-qPCR assays presented efficiencies between 99.6% and 105.2%.

Two of the 6 circRNAs showed significant differences between the two sperm motility groups (Figure 3.B). These 2 circRNAs were ssc_circ_1458 from *LRBA* (P-value: 0.049) and ssc_circ_1321 from *PAPOLA* (P-value: 0.035). A third circRNA, ssc_circ_1132, from *LIN7A*, showed significant differences between the two motility groups (P-value: 0.008) (Figure 3.B) but in the opposite direction than expected according to the RNA-seq data and was consequently considered as not validated due to inconclusive results. The other 3 circRNAs, ssc_circ_1101 from *KHDRBS3*, ssc_circ_0437 from *ULK4* and ssc_circ_1061 from *ZNHIT6* did not present significant differences between the 2 groups (Figure 3.B).

## Discussion

In this study, we have profiled for the first time, the circRNA repertoire of mature spermatozoa in a mammalian species and its association with sperm motility as a parameter of semen quality. These results provide further evidence and expand the relevance of spermatoza RNAs in sperm biology and quality. circRNAs have been reported as promising potential prognostic and diagnostic biomarkers for human health including, among others, cancer, diabetes, cardiovascular diseases or pre-eclampsia [reviewed in 30]. Here, we provide novel data supporting the potential of circRNAs as biomarkers for the male’s reproductive function as reflected by association with sperm motility parameters.

We have identified nearly 1,600 boar sperm circRNAs that are robustly present in most samples (at least 30). The sperm circRNA repertoire presented similar genomic circular characteristics such as the proportion of genomic overlap with protein coding or intergenic regions, among others, the number of exons and their length (Figure 1), when compared to data previously reported in other tissues from swine [28], human [44] or rat [41]. Nonetheless, the list of sperm circRNAs showed a modest overlap with other tissues and species (Table 3). Not surprisingly, the largest overlap was with porcine testes (11.6%) probably due to the fact that the spermatogenic lineage including spermatozoa is contained in the male gonads. The proportion of pig sperm circRNAs shared across swine tissues was in general low when compared to the rest of the *Sus scrofa* libraries (Supplementary Table 4). Although this comparative analysis may be partly influenced by technical differences involving the processing of the tissues and the data between the studies, our results suggest that sperm present high tissue-specificity, with circa 80% of the circRNAs being present only in sperm, considerably higher than testes [28], with approximately 65% testes-specific. On the opposite direction, there was a larger coincidence between the catalogue of around 6,660 porcine circRNAs [28, 29] and the swine sperm circRNAome (4.9%), than between the later and the human archive with over 90,000 circRNAs [36] (0.4%) (Table 3). This 12x fold difference indicates that circRNAs are not only tissue specific but have also a strong species-specific component.

We investigated the functional relevance of sperm circRNAs, under the hypothesis that their function is associated to the host gene. Six (*CEP63, ATP6V0A2, PPA2, PAIP2* and *PAXIP1*) of the 15 coding genes providing the top 20 most abundant circRNAs (Table 1) have been directly implicated, to sperm related traits and male fertility. *CEP63* is engaged in microtubule organization, and mice knockout studies revealed its essential role in male fertility [45]. *ATP6V0A2* proteins and transcripts are down-regulated in infertile men [46]. *PPA2* is located in the mitochondrial membrane and might be involved in the production of ATP, control of molecular processes linked to the launching of sperm capacitation and sperm motility [47–49]. *PAIP2* is linked to male sperm maturation and fertility [50] and *PAXIP1* is associated with developmental arrest of spermatocytes, testicular atrophy, and infertility in knockout mice [51]. We identified 12 circRNA from hotspot genes producing five or more circRNAs each (Table 2). Some of the hotspot genes were related to sperm function and fertility. *TESK2* may play a role in early stages of spermatogenesis [52]; *PTK2*, which is essential for a embryo development [53]; *SPATA19*, a gene that is critical for sperm mitochondrial function in relation to sperm motility and fertilization ability [54], and *SLC5A10*, which protein products may act as water channels in spermatozoa [55].

The ontology enrichment analysis of the genes harboring circRNAs pointed towards epigenetic related functions, which are essential in all cell types including chromatin condensation in sperm and the reprogramming of gene expression upon egg fertilization and during embryo development (Supplementary Table 2). Gene enrichment analysis also signaled towards spermatogenesis and developmental processes, the later also implicating the embryo development related genes, *DHX36, IPMK, RICTOR, CDC73* and *ANGPT1* which were hosting some of the circRNAs found in our analysis (Supplementary Table 1). These functions are in line with the gene ontologies highlighted in studies analyzing sperm mRNAs [34, 35, 56], thereby providing further basis for the hypothesis that circRNAs may exert their function (regulation of sperm quality and motility in our study) by controlling their cognate linear mRNA.

A previous study in rat testes identified a circRNA age-dependent dynamic pattern of expression and suggested a relation between their abundance and function with the male’s sexual maturity and spermatogenesis [41]. For this reason, we sought to investigate whether, like in testes, sperm circRNAs - in boar at least - also accumulate through age. Our data highlighted that there is no association between the mature sperm circRNAs and the boar’s age. Cells with high proliferation rates, seem to accumulate less circRNAs, possibly due to passive thinning out during proliferation [57]. Spermatogenesis is a process that occurs throughout the male’s lifetime in which spermatogonial stem cells (SSCs) undergo continuous cell division and finally differentiate to ultimately become spermatozoon [58]. Thus, the fact SSCs undergo continuous self-renewal and keep proliferating through the male’s lifetime may impede the accumulation of circRNAs in these cells and thus explain why no age-dependent pattern of circRNA abundance was found in the sperm of pigs with different ages. It is plausible that due to the continuous production and differentiation of the male germ cell, the sperm circRNAome does not mimic the reproductive performance of the boar, at least once the number of germ cells have stabilized which occurs at the age of 7 months old in pigs [59] and until the boar is senile. These results infer that stability of mature sperm circRNAs might be constant for a period in normozoospermic males and provides further evidence for circRNAs as a noninvasive diagnosis tool for male reproductive diseases.

To elucidate the functional relevance of circRNAs as miRNA sponges we built an interaction network (Figure 2). We combined benchwork data of circRNA and miRNA abundances (small and total RNA-seq) and *in silico* searches of miRNA target sequences in circRNAs with the aim to increase the reliability of the circRNA-miRNA relationship predictions. This network with 70 interactions, contained some interesting circRNAs and miRNAs in relation to semen quality and male fertility. Remarkably, two circRNAs, ssc_circ_0954 and ssc_circ_1454, presented, each, 4 different miRNA target sites. The first, ssc_circ_0954 arises from *DCDC2C*, a gene identified in the human’s sperm flagellum end-piece with a suggested role on microtubule dynamics by acting as a depolymerization/polymerization balancing system [60]. This circRNA regulated among others, miR-361-3p which has been found dysregulated in subfertile men [61], miR-423-5p, altered in oligozoospermic men [62] and miR-28-5p, dysregulated in normozoospermic infertile individuals [18]. The other cirRNA, ssc_circ_1454, is transcribed from *MTHFD2L*, a mitochondrial isozyme from the folate cycle metabolic pathway [63], a vitamin that has been also related to semen quality and fertility in men [64]. One of the miRNAs targeted by ssc_circ_1454 was miR-16, which was found dysregulated in subfertile men [61]. The network also showed 9 circRNAs that may be regulating miR-28, a miRNA (miR-28-5p) that is dysregulated in normozoospermic infertile individuals [18]. The potential miR-28 regulators include ssc_circ_1370, arisen from *FAM92A*, whose protein may play a role in ciliogenesis [65], implicated in the formation of the sperm flagella and ssc_circ_0002 from *WDR7*, which is associated to cattle sperm quality [66]. Likewise, 9 circRNAs were predicted to regulate miR-26a, which has been in turn, linked to VCL, VSL and VAP motility parameters in swine [67]. These miR-26a regulatory edges implicate ssc_circ_0361 from *ACTL6A*, crucial for embryo development [68] and ssc_circ_1352 from *CAGE1*, an acrosomal protein with proposed roles in fertility [69]. Other interesting interactions included ssc_circ_0345 from the hotspot gene *SLC5A10* (sodium-dependent mannose and fructose transporter) (Table 2), which regulated miR-423-5p, a miRNA that was found to be upregulated in oligozoospermic semen [62] and let-7c, which is altered in severe aasthenozoozpermia patients [70]. Altogether, the network involves key genes and miRNAs for male fertility thereby suggesting that circRNAs play a functional role in the male reproductive ability.

The potential role of circRNAs in male reproductive traits is further substantiated by the associations between sperm motility and the abundance of some circRNAs in our study. At least 20 of the circRNA host genes implicated in the phenotypic correlations have been previously linked to sperm biology or male fertility (Supplementary Table 3). For example, ssc_circ_0823, which correlated with VCL (p-val = 0.009), is a circRNAs hosted by *CAMK4*, a gene that has been implicated in sperm motility in humans [71]. ssc_circ_0780, correlated with the percentage of motile cells (p-val = 0.041) from *LRGUK*, a gene required for sperm assembly including the growth of the axonome, a structure that is necessary for the flagellar beating in sperm [72]. We successfully tested by RT-qPCR 6 of the 179 circRNAs that showed RNA-seq based significant correlations between their abundance and sperm motility. The results allowed us to validate the correlation with motility for 2 of these 6 circRNAs, ssc_circ_1458 from *LRBA* and ssc_circ_1321 from *PAPOLA*. Both were significantly down-regulated in the ejaculates with low sperm motility values (Figure 3.B). *LRBA* is a gene involved in coupling signal transduction and vesicle trafficking but no link with sperm function or fertility has been made thus far. This gene has been associated to immune-related disorders in humans. ssc_circ_1321 from *PAPOLA*, has a human ortholog (human circRNA: hsa_circ_0033126) and the host gene is implicated in RNA and ATP binding. Thus, none of the two host genes have been previously related to sperm function or fertility. Moreover, these 2 circRNAs are not included in the circRNA:miRNA interaction network. However, the association is clear and confirmed by RT-qPCR and we cannot exclude unidentified relevant functions on sperm motility for these two genes. Two other circRNAs were also of high interest and tested as they were within the top 20 most abundant (Table 1) and correlated with sperm motility (Supplementary Table 3). They were ssc_circ_0839 from *PAIP2*, with crucial roles in spermatogenesis [50] that did not amplify, and ssc_circ_1101 from *KHDRBS3*, found highly abundant in mice testes [73], which did not present significant differences in RT-qPCR levels between motility groups (Figure 3.B).

Interestingly, some circRNAs popped up as relevant in more than one of the analysis carried through the study. For example, ssc_circ_1532 from the hotspot gene *SPATA19* (Table 2), related to sperm motility and fertility [54], was suggested to regulate miR-99a according to the network analysis (Figure 2). miR-99b has been found deregulated in low motile sperm fractions in bull [19] and in subfertile men [61]. Another circRNA, ssc_circ_1219 from *OSBPL9*, involved in male reproduction [74], displayed abundance correlation with VCL (P-value: 0.04) (Supplementary Table 3) and was identified as a potential target of miR-101 (Figure 2), a miRNA (miR-101-3p) that was altered in asthenozoospermia men [70].

Remarkably, 4 (*DENND1B, PTK2, SLC5A10* and *CAMSAP1*) of the 12 hotspot genes hosted a circRNA with significant abundance correlation with sperm motility. Thus, one third of the hotspot genes included circRNAs correlated with motility whilst only 14.9% of the genes (147) of the 984 genes hosting the 1,598 circRNAs were correlated to motility. Noteworthy, the 12 circRNA hotspot genes were not hotspots in the other porcine tissues analyzed [28, 29] (data not shown). Altogether, this indicates that circRNAs hotspot genes may have relevant tissue-specific functions.

CircRNAs are acknowledged to be more stable than mRNA. Our data shows, at different levels, that there is a detectable population of circRNAs in the boar sperm that is related to relevant functions in sperm and fertility. Moreover, these circRNAs are stable across age. For these reasons, our results indicate that circRNAs hold a potential as biomarkers for sperm motility and potentially, fertility outcomes [7, 8, 10–14, 75]. This potential should be further explored in swine, as well as in other domestic animals and in human fertility clinics.

In conclusion, our study is the first to characterize the spermatozoa circRNAs repertoire in an animal species. We have provided a comprehensive view of the boar sperm circRNAome and their correlation with sperm motility parameters. Our findings involving sperm motility may spur novel research on male fertility in both human medicine and in animal breeding.

## Material and methods

### Sperm collection, phenotyping and library preparation

Ejaculates from 300 Pietrain boars were collected using the hand glove method by trained professionals at commercial farms. Fresh sperm motility traits were assessed with the CASA system (Integrated Sperm Analysis System V1.0; Proiser, Valencia, Spain). In this study we analyzed sperm motility parameters, including the total percentage of motile cells, VCL (µm/s), VSL (µm/s) and VAP (µm/s). The average percentage of motile cells was 75.1 with a standard deviation (SD) of 18.3, VCL (mean: 45.1; SD: 12.6), VSL (mean: 27.0; SD: 8.3) and VAP (mean: 34.2; SD: 10.5). Phenotypes were corrected for the fixed variables: farm (1, 2, 3), age (1, 2, 3) and season and year (Autumn 2014, 2015 and 2016; Winter 2015, 2016 and 2017; Spring 2015 and 2016; Summer 2015) using the R function “lm” [76]. Ejaculates were purified to remove somatic cells and immature sperm cells and purified sperm was stored at −96°C with Trizol^®^ as described by Gòdia et al. [77]. RNA was extracted from purified sperm cells, treated with TURBO DNA-free™ Kit (Invitrogen; Carlsbad, USA) and quantified using Qubit™ RNA HS Assay kit (Invitrogen; Carlsbad, USA). The RNA yield of these samples averaged 2.2 fg per sperm cells (the range was between 0.8 and 3.7 fg). We assessed RNA integrity with the 2100 Bioanalyzer using the Agilent RNA 6000 Pico kit (Agilent Technologies; Santa Clara, USA). All samples presented RNA Integrity Number (RIN) below 2.5, which indicates the absence of intact RNA from somatic cell origin. We then performed RT-qPCR assays for *PRM1* and *PTPRC* mRNAs as well as for intergenic/genomic DNA to verify that all the samples were free from RNA from somatic cells and from genomic DNA contamination [77].

40 sperm RNA samples were subjected to total RNA-seq. 34 of these samples were also used for small RNA-seq. For total RNA-seq libraries, ribosomal RNA (rRNA) was depleted with the Ribo-Zero Gold rRNA Removal Kit (Illumina) and libraries were constructed with the SMARTer Universal Low Input RNA library Prep kit (Clontech). Resulting libraries were sequenced in a HiSeq2500 system (Illumina) to generate 75 bp long paired-end reads. For small non coding RNA-seq libraries, extracted RNA without rRNA depletion was directly subjected to library preparation with the NEBNext Small RNA Library Prep Set kit (New England Biolabs) and sequenced on a HiSeq2000 (Illumina) to generate 50 bp single-end reads.

### RNA-seq and bioinformatic analysis

For the total RNA fraction, we obtained, in average, 20.4 million paired-end reads per sample. Raw reads were then filtered by removing adaptor sequences and low-quality reads with Trimmomatic v.0.36 [78]. In average, 98.5% of these reads passed the quality control filters. The identification of circRNAs was carried on these reads with the find_circ pipeline [26] with stricter filter stringency. To reduce the false positive rate in the discovery of circRNAs, we selected circular splice transcripts with at least two unique supporting reads in the anchor segment and with Phred quality scores of 35 or more. Moreover, only circRNAs predicted in at least 30 samples were kept. The RNA abundance of the predicted circRNAs were normalized as the number of back-splice junction spanning reads by its sequencing depth, as counts per million (CPM). The functional regions of circRNAs were identified based on their co-location with genomic features (e.g. exon, 3’UTR, 5’UTR, etc) from the Ensembl database (release 91) with BEDtools [79]. Our catalogue of boar sperm circRNAs was contrasted with other publically available porcine circRNA databases including heart, liver, spleen, lung, kidney, ovary, testis, skeletal muscle, fat and fetal brains [28, 29]. We also queried several human tissues (including several cell lines, brain sections placenta, muscle, fat, umbilical cord, atrium, decidua and plasma) [26, 37–40] and murine (cell lines and brain sections) [26, 37] available at the circBase database [36]. Genomic coordinates from the human and mouse circRNAs were liftover to Sscrofa11.1 using the UCSC liftover tool [80].

For the small RNA-seq analysis, we obtained an average of 7.3 million reads per sample. Trimming of adaptors and low quality bases was performed with Cutadapt v1.0 [81]. 99.2% of these reads were of high quality and were thus used for downstream analysis. The mapping of sncRNAs was performed with the sRNAtoolbox v.6.17 [82] with default settings and providing miRBase [83] release 21 as library dataset. Multi-adjusted read counts were then normalized by sequencing depth as CPM. We only considered the miRNAs that were detected > 1 CPM in all the samples.

### circRNAs-miRNAs network visualization

For a functional annotation of the circRNA and miRNA interactions in a systems biology context, we carried a Partial Correlation with Information Theory (PCIT) analysis [84] using the combined list of circRNAs and miRNAs after normalization of abundance levels with log2. We further assessed circRNAs-miRNAs co-abundance interactions using miRanda [85] v.3.3a. Shared correlations were visualized with Cytoscape v.3.7.0 [86].

### Genome Ontology analysis and correlation with sperm motility parameters

GO analysis was carried with PANTHER v.13.1 [87] with the overrepresentation test and p-values corrected with FDR. Annotation Data Set was “GO biological process complete”. Pearson correlation was used to determine associations between circRNA abundance levels and sperm motility parameters. P-values < 0.05 were considered statistically significant.

### Validation of circRNAs and reverse transcription quantitative PCR (RT-qPCR)

circRNAs were validated by Sanger Sequencing and quantified by RT-qPCR using divergent primers. Primers were designed as in [88] using the Primer Express software (Applied Biosystems). Primer sequences are shown in Supplementary Table 5. For cDNA synthesis, 5 µl of RNA were reverse transcribed using the High Capacity cDNA Reverse Transcription kit in a final volume of 50 μL (Applied Biosystems; Waltham, USA) following the manufacturer’s protocol. circRNAs were amplified and visualized in 3% high resolution agarose gel electrophoresis and confirmed by Sanger Sequencing.

The abundance level of 6 circRNAs, correlated to sperm motility parameters in the RNA-seq study, was analyzed by RT-qPCR in 36 samples, none of them included in the RNA-seq. These 36 samples belong to two groups with extreme and divergent values for sperm motility from a bank of 300 ejaculates with phenotypic records. Quantitative PCR reactions were performed in triplicate in 15 μL final volume including 7.5 μL SYBR Select Master Mix (Life Technologies - Thermo Fisher Scientific), 300 nM of each primer and 3.75 μL of cDNA 1:4 diluted on a QuantStudio 12K Flex Real-Time PCR System (Applied Biosystems). To evaluate the efficiency of the RT-qPCR assays, standard curves with 6 serial dilutions from a pool of sperm cDNA were generated. Thermal profile was set as follows: 50°C for 2 min, 95°C for 10 min and 40 cycles at 95°C for 15 sec and 60°C for 60 sec. Moreover, a melting profile (95°C for 15 sec, 60°C for 15 sec and a gradual increment of temperature with a ramp rate of 1% up to 95°C) was programmed at the end of the RT-qPCR to assess the specificity of the reactions. The genes *ISYNA2* and *GRP137* were selected as endogenous controls following the stability values after a GeNorm pilot experiment. Their stability was determined considering a GeNorm M value < 0.5. Relative expression values were calculated using the ThermoFisher Cloud software (Applied Biosystems) applying the 2^−ΔΔCt^ method. The same software was used to compare the biological groups. Significance was set at a P-value < 0.05.

## Author contributions

MG, AS, and AlC conceived and designed the experiments. MR and JR-G carried the phenotypic analysis. MG performed sperm purifications. MG and MR carried the RNA extractions. AnC designed and carried the RT-qPCR and their analyses. AnC and BC performed Sanger Sequencing validation. MG made the bioinformatics, statistic analysis and analyzed the data. MG and AlC wrote the manuscript. All authors discussed the data and read and approved the contents of the manuscript.

## Availability of data

The datasets generated and/or analysed during the current study are available at NCBI’s BioProject PRJNA520978.

---

**Supplementary Figure 1.** Validation of the amplified set of circRNAs by agarose-gel. **A.** Amplification of the 2 randomly selected circRNAs. Two different primer sets for ssc_circ_1141 from *PTGES3* were tested (primer pair a in lane 2 and b in lanes 3 and 4) and ssc_circ_0670 from *BAZ2B* (lane 5). **B.** Validation of the circRNAs correlated to sperm motility parameters: ssc_circ_1321 from *PAPOLA* (*ENSSSCG00000002505)*, ssc_circ_1458 from *LRBA*, ssc_circ_0437 from *ULK4*, two different primer sets for ssc_circ_1061 from *ZNHIT6* primer pair a and b, ssc_circ_1132 from *LIN7A*, two different primer sets for ssc_circ_1101 from *KHDRBS3*, primer pair a (which resulted in amplification of two splicing forms and was excluded) and b, and ssc_circ_0118 from *PDE10A* that resulted in amplification of two splicing forms (and excluded from further analysis).

**Supplementary Figure 2.** Sanger sequencing validation of the set of circRNAs. **A.** ssc_circ_1141 from *PTGES3*. **B**. ssc_circ_0670 from *BAZ2B*. **C.** ssc_circ_0437 from *ULK4*. **D.** ssc_circ_1061 from *ZNHIT6*. **E.** ssc_circ_1101 from *KHDRBS3*.

## Supplementary Tables

**Supplementary Table 1.** List of the 1,598 circRNAs identified in sperm with coordinates of the exonic host gene and the mean and Standard Deviation (SD) of the RNA abundances (in CPM) in the 40 samples.

**Supplementary Table 2.** Gene Ontology analysis and FDR value of the circRNA host genes.

**Supplementary Table 3.** Correlation of circRNAs with sperm motility parameters. Including the exonic host gene, if tested for validation and article reference when the host gene has been associated to sperm biology or male fertility. MT: total percentage of motile cells; VCL: curvilinear velocity; VSL: straight line velocity; VAP: velocity of the sperm cells; ns: not significant.

**Supplementary Table 4.** Comparison across swine circRNAs tissues.

**Supplementary Table 5.** CircRNA and reference genes primers designed.

