## Supplementary figures and images for "Identification of circular RNAs in porcine sperm and their relation to sperm motility"

### Supplementary Figure 1

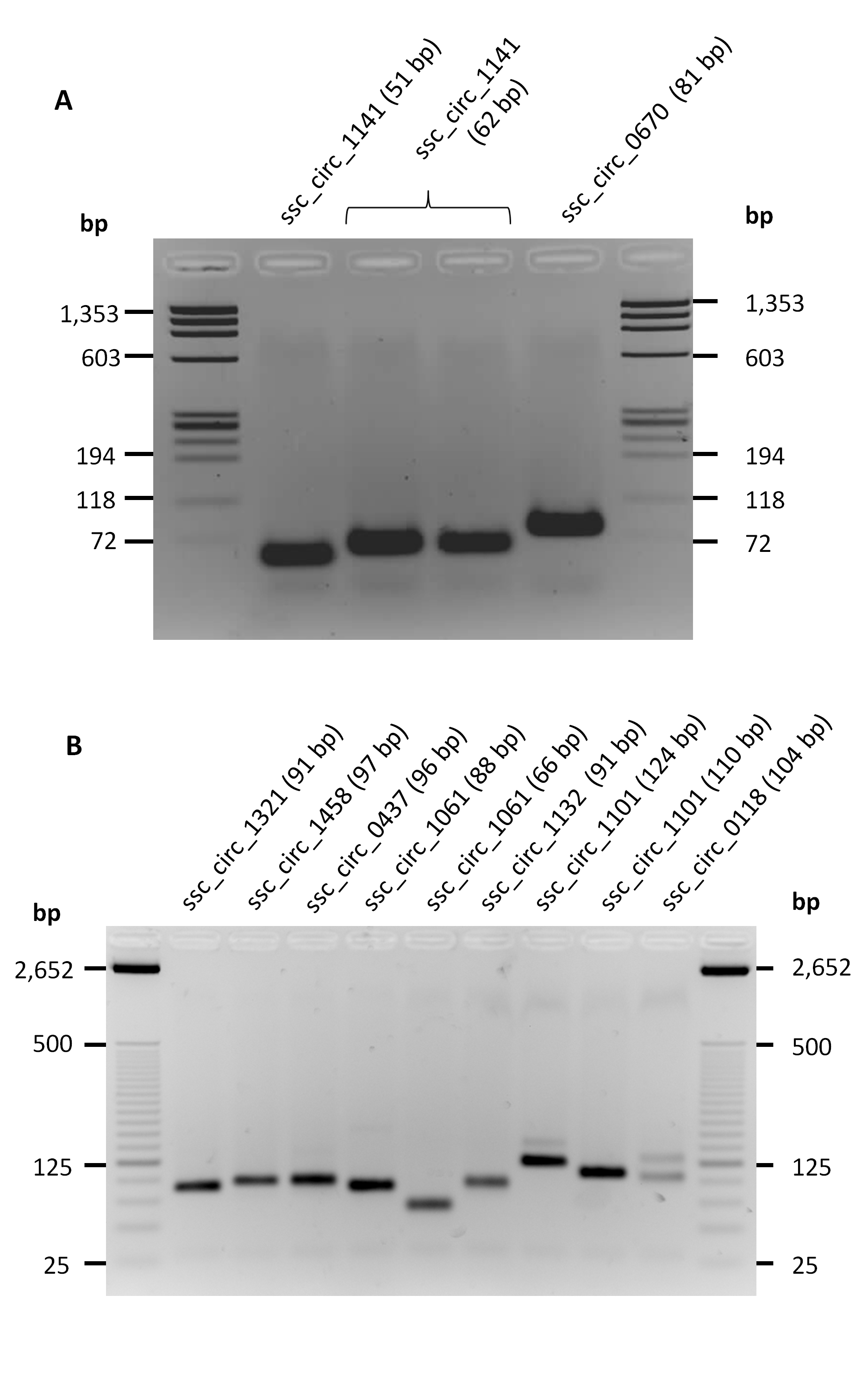

### Supplementary Figure 2

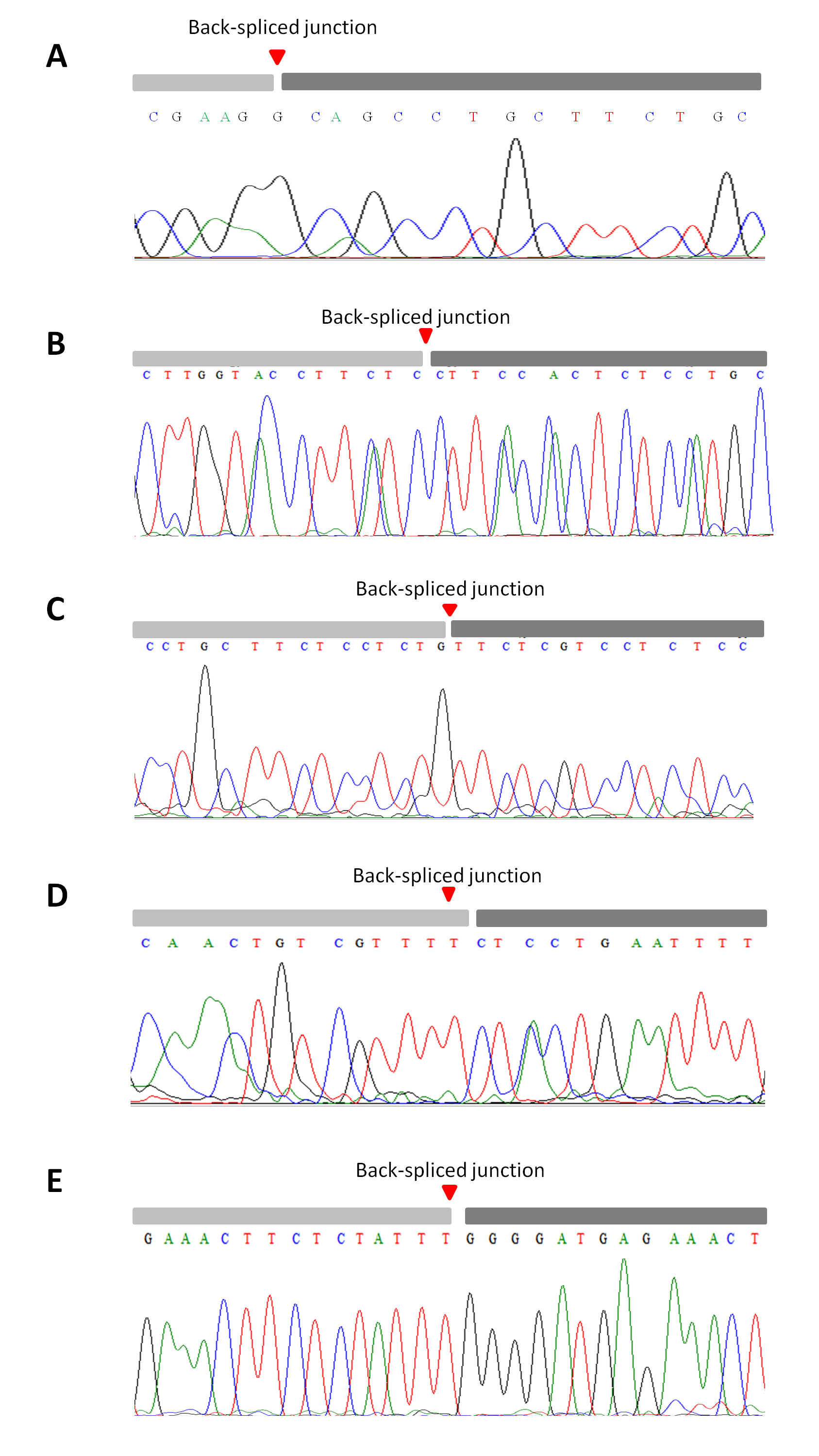
